# Lightsheet Optical Tweezer (LOT) for Optical Manipulation of Microscopic Particles and Live Cells

**DOI:** 10.1101/2021.07.05.451038

**Authors:** Partha Pratim Mondal, Ankur Singh, Prakash Joshi, Jigmi Basumatary, Neptune Baro

## Abstract

A light sheet based optical tweezer (LOT) is developed to trap microscopic dielectric particles and live HeLa cells. The technique requires the generation of a tightly focussed diffraction-limited light sheet which is realized by a combination of cylindrical lens and high NA objective lens. The field pattern generated at the geometrical focus is a tightly focussed line (along *x*-axis) perpendicular to the beam propagation direction (*z*-axis). Spherical beads undergoing Brownian motion in the solution are trapped by the gradient potential, and the time (to reach trap-center under the influence of gradient potential) is estimated from the fast CMOS camera (operating at 230*frames*/*sec*). High-speed imaging of beads at varying laser power shows a steady increase in the stiffness of LOT with a maximum of 0.00118 *pN*/*nm* at 52.5 *mW*. This is an order less than traditional optical point-traps. The trapped beads displayed free movement along the light-sheet axis (*x*-axis), exhibiting a single degree of freedom. Subsequently, LOT is used to optically trap and pattern dielectric beads and HeLa cells in a line. We could successfully pattern 8 dielectric beads and 3 HeLa cells in a straight line. We anticipate that LOT can be used to study the 1D-physics of microscopic particles and help understand the patterned growth of live cells.

---

Optical tweezers (OTs) are known to play key roles in understanding molecular forces (or torques) and mechanical properties of proteins (DNA, RNA, etc.). In addition, this techniques has applications in optical sorting (sorting cells [2] [3], sorting colloidal spheres [1]), and single-molecule biophysics [4] [5]. The delicate force exerted by radiation has consequences in precision measurement, determination of intermolecular forces, and short-distance interactions in sub-piconewton range.

In the year 1970, Arthur Askhin was the first to show that micron-sized latex spheres suspended in water can be manipulated [6]. Subsequently, the first demonstration of single-beam optical tweezers was carried out, and successful trapping of bacteria and red blood cells was realized [7] [8]. The basic physics of optical tweezers revolves around the fact that light carries linear and angular momentum, and this can be harvested to manipulate microscopic particles both inside and outside live cells. The forces that compete and need to be balanced for a stable trap are gradient and scattering forces. For his pioneering work on optical tweezers, A. Askhin received nobel prize in the 2018 [9]. The field is growing at a rapid pace and finding new applications across the disciplines of science and engineering.

Light focused by spherical optics (such as biconvex lens or objectives) produces point focus with maximum intensity at the centre. This gives rise to a well-defined point traps when focussed by light with a Gaussian beam profile (such as laser). Both sub-micron and micron sized particle can be trapped and manipulated using point traps. This is different for cylindrical lens system that focus light to a line rather than point. As a result, the particle trapped in line-focus has one-degree of freedom but restricted in the other two directions. Historically, light sheet was first generated by Siedentopf and Zsigmondy in 1903 to observe gold particles [10]. Later on, a modernized version of light sheet microscopy was built using cylindrical lens in 1993 by Voie et al. [11]. Subsequently, light sheet was diversified by Stelzer and Keller for biological imaging [12] [13] [14]. The technique has seen applications in diverse fields ranging from biological sciences (biological imaging [15] [16] [17] [18] [19] [20], iLIFE imaging cytometry [38] [21]) to physical sciences (nanolithography [24] [25], optics [22] [23]). The last decade has seen an emergence of several important variants of light-sheet systems. Some of these include thin light-sheet microscopy [26], ultramicroscopy [27], objective coupled planar illumination microscopy (OCPI) [28], confocal light-sheet microscopy [29][30], multiple light-sheet microscopy [31], dual-inverted selective-plane illumination microscopy (diSPIM), [32], light-sheet theta microscopy (LSTM) [33], open-top light-sheet (OTLS) [34][35] and lattice light sheet microscopy [36], LVLSM [37] and IML-SPIM [17]. These variants offer many useful configuration for generating light sheets that can be used for demanding applications.

In this letter, we propose and develop lightsheet optical tweezers (LOT) for trapping microscopic objects. This is accomplished by generating a diffraction-limited lightsheet using a combination of cylindrical lens and a high NA objective lens. A stable and confined optical trap is realized, and the same is used to demonstrate the trapping of dielectric beads and live HeLa cells.

The theory of line-traps (produced by a light sheet) is similar to that of point focus, with the exception that particle has complete freedom along one dimension. Unlike point-traps that are better understood in the cylindrical coordinate system (*r, z*, with *r* being the lateral/radial plane and *z* being the beam propagation direction), LOT is better understood in a rectangular coordinate system (*x, y, z*) as shown in Fig. 1. The diffraction-limited light sheet is taken along xz-plane with *x* and *z* as the lightsheet axis and beam propagation direction, respectively. From Fig. 1, the cylindrical lens focus light on a line extending along *y*-axis (see, yellow oval just before the back-aperture of the objective lens in Fig. 1B). The objective lens is placed at the focus of a cylindrical lens. The field at the back-aperture of objective lens undergoes Fourier transform, forming a diffraction-limited line at its focus (see, yellow oval along *x*-axis at the focus of objective lens in Fig. 1B). In general, two cases arise: (1) the particle is much smaller than the wavelength of light (Rayleigh regime), and (2) the particle is larger than the wavelength of light (Geometric regime). We briefly discuss both cases.

**FIG. 1:**
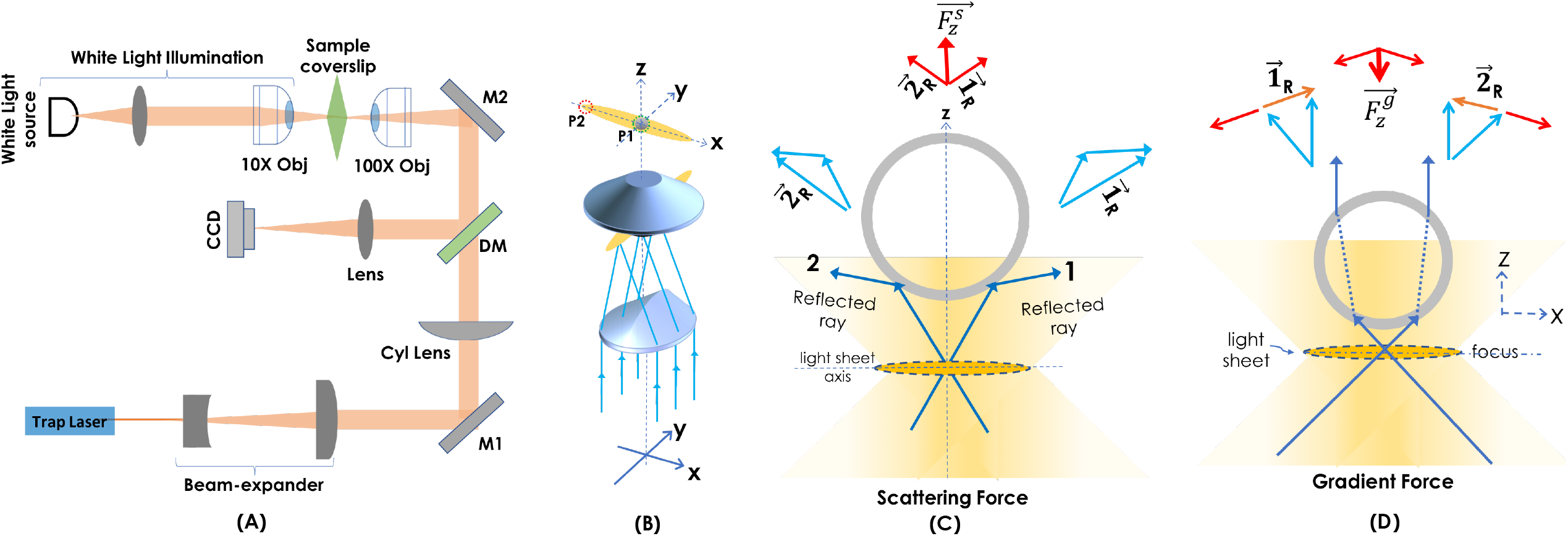
(A) Schematic diagram of the developed lightsheet optical tweezer (LOT) system. (B) The combination of cylindrical lens and high NA objective lens for generating diffraction-limited light sheet. (C) The resultant scattering force (due to the reflection of light) acting on the micro-particle along +*z*-axis. (D) The gradient force (purely due to intensity-difference) acts inwards (towards lightsheet axis) on the particle, i.e., along the –*z*-axis. The force diagrams shown with blue and red arrows indicate elemental and resultant forces, respectively.

In the Rayleigh regime, we consider a dielectric particle in a laser beam and that the size of particle is much smaller than wavelength of light. In this regime, the particle behaves as a Rayleigh scatterer, and its reaction to the light field, 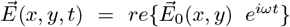 is determined by its polarizability (*α*). The electric field on the particle induces a dipole moment, 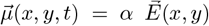, and the energy of induced dipole is, 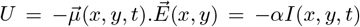, where, 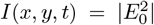. This immediately states that the energy of dipole assumes minimum at a position where the intensity is maximum, resulting in a well-defined trap at (*x,y*). This rules out optical trapping for a uniformly intense light source, and thus a non-uniform intensity profile such as those for Gaussian beam is desired. In the lateral case of beam with Gaussian intensity profile, the dipole (particle) is expected to experience a force towards higher intensity i.e, the beam centre. The corresponding force along *y* is given by, 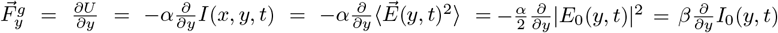, where, 〈…〉 is the time average, and 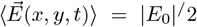. Note that, the variation of intensity along *x* is negligible and does not change appreciably, except at the far ends. Hence, 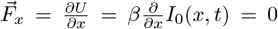 along *x*. In the above expression, we have absorbed the refractive index and permittivity in a single parameter, *β* = *α*/2*cn∊*_0_, where *n* is the refractive index of the particle and *∊*_0_ is the permittivity of vacuum. Accordingly, the particle is trapped when the polarizability of particle is greater than the surrounding media. In the geometric regime, where the particle size is larger than the wavelength of light such as dielectric beads, ray-optics can be employed to understand forces acting on the bead / particle. Classically, force on the particle can be defined as the rate of change of momentum, 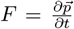, where 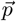 is the momentum of the particle. The conservation of momentum necessitates the exchange of momentum between light and the particle. However, the off-focal beads experiences a net force towards the trap-center (high intensity region) due to gradient force as explained by the force diagram in Fig. 1D. A similar force but in opposite direction appears when the particle is on the other side of light sheet axis.

Now, let us determine the role of scattering force *F_s_*. Primarily, the scattering occurs due to reflection of light, and the scattering force in Rayleigh regime can be expressed as, 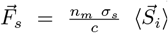, where, *σ_s_* is the cross-section of particle. This means that scattering forces are directly proportional to the cross-section of particle. So, large particles experience greater scattering force. Fig. 1C shows the resultant scattering force 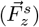 due to the reflection of light at the bead surface. This points in the direction of Poynting vector 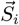 which is also the direction of beam propagation (z) and has a unit of energy per unit area per unit time. So, scattering force has the direction of Poynting vector. Accordingly, scattering (due to reflected light at the bead surface) result in momentum transfer between light and particle that tends to push the particle out of focus with force, 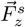 (see, Fig. 1C). So, the gradient force with its maximum at the centre leads to a stable trap both along the beam propagation direction (*z*-axis). This is similar along *y*-axis as well. In LOT, the bead is free to move along *x*-axis due to negligible intensity gradient. Overall, the condition for three-dimensional stable trap is realized when the gradient potential overcomes the other forces (radiation pressure or scattering force, buoyant force, and the forces due to Brownian motion and gravity).

The next logical step in the development of LOT is to determine the trap-stiffness. We used spherical transparent beads as the sample to estimate trap-stiffness. Within the influence of gradient force, the particles/beads undergoing Brownian motion are attracted towards the trap center. To a good approximation, an optical trap behaves like a harmonic potential, and it is able to exert a restoring force. Specifically, near the trap center, the force can be approximately modeled by Hooke’s law, and the restoring/gradient force is given by *V*(*x*) = –*kx*, where *k* is the trap stiffness (N/m) and *x* is the travel distance/displacement from the trap origin. The second force acting on the particle is viscous drag force. Assuming spherical particle (beads), the particle moving through the fluid experiences a viscous force of, *F_vis_* = −6*πηr_b_v*. For simplicity and calculating approximate trap-stiffness, we neglect the effect of gravity on the bead so we can ignore forces due to weight and buoyancy. Thus, the motion of bead is governed by these two forces (gradient and viscous/drag force), which are opposite. Balancing optical (gradient) forces with drag force produces [39] [40], –*kx* = −6*πηr_b_v* ⟹ *k* = 6*πηr_b_*(*v/x*) ⟹ *k* = 6*πηr_b_*/*t*, where *x* = *vt* and, *v* is the dragging velocity, *x* is the displacement and *t* is the time, *η* is the medium viscosity, and *r_b_* = *d/2* is the bead radius. We will use this relation to determine the trap-stiffness.

The schematic diagram of the developed optical tweezer (LOT) is shown in Fig. 1A. The picture of actual optical tweezer is shown in the **Supplementary Fig. S1**, and related description can be found in **Supplementary material**. A trap laser light of wavelength 1064 *nm* is used to trap dielectric silica beads (Thorlabs, USA). The laser beam is expanded 3 times by the beam expander so as to fill the back-aperture of cylindrical lens (Cyl Lens, *f* = 150 *mm*). The lens focusses light along *y*-axis on to the back-aperture of 100X objective lens (Olympus, 100X, 1.25 NA), without any change along *x*-axis. The line focus at the back-aperture of objective lens gives rise to an orthogonal diffraction-limited line-focus (see, Fig. 1B). A separate illumination sub-system is integrated for visualizing the specimen (beads and cells in solution). The illuminator consists of a white light source a condenser lens and a low NA objective lens (Olympus, 10X 0.25 NA). The lens illuminates a larger field-of-view (FOV) of the sample plane, and the transmitted light is collected by the 100X objective lens. The light then pass through the dichroic mirror (DM) to the tube lens which focus it to the fast CMOS camera (Gazelle, Pointgray, USA). A schematic of the key optical element for generating light sheet trap is shown in Fig. 1B. Note that, the light sheet generated by the cylindrical lens is projected on to the back-aperture of high NA objective lens that give rise to a diffraction-limited light sheet at the focus. The line focus is formed along the x-axis and orthogonal to propagating direction (along *z*-axis). The resultant field and trap geometry is shown in Fig. 1. Two major forces (gradient force and scattering force) acting on a spherical bead are shown in Fig. 1(C,D). The scattering force 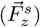 on the particle is towards the beam-propagation direction (z-axis outwards) that pushes the particle away from the light sheet centre (see, red arrow). The corresponding vector forces acting on the particle is shown in Fig. 1C. On the other hand, the gradient force is primarily due to refraction and exerts a restoring force on the particle when it is away from the centre (lightsheet axis). So, the gradient force 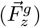 pushes the particle towards high intensity which is the centre of trap (see, red arrow). This is explained based on the vector force diagram shown in Fig. 1D. Unlike point-traps where particle experience gradient forces radially inwards (along 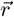, LOT involves gradient force along *x* and *y* directions only. This leaves the particle free to move along x-axis without experiencing any net force.

Trapping micro-particles require a strong, stable and confined optical field (system PSF). This can be generated by focussing high intensity light by a high NA objective lens. The schematic diagram of the key optical component along with the formation of light sheet is shown in Fig. 2A. The field is shown at discrete values of *z* in the specimen. Fig. 2B shows the actual field recorded by the camera (in the reflected mode) for a combination of cylindrical lens (*f* = 150 *mm*) and high NA objective lens (100*X*, 1.25*NA*). Visually, the field displays optical aberration in the specimen medium which is predominantly due to multiple reflections and medium inhomogeneity. Alongside the intensity plots are also displayed (see, Fig. 2C). During experimentation, we noted that trapping is efficient for slightly saturated light sheet. Gaussian function is fit to the data to determine the dimension of light sheet. The size of light sheet is estimated to be 14.23 *μm* along *x*-axis and has a thickness of 0.87 *μm*.

**FIG. 2:**
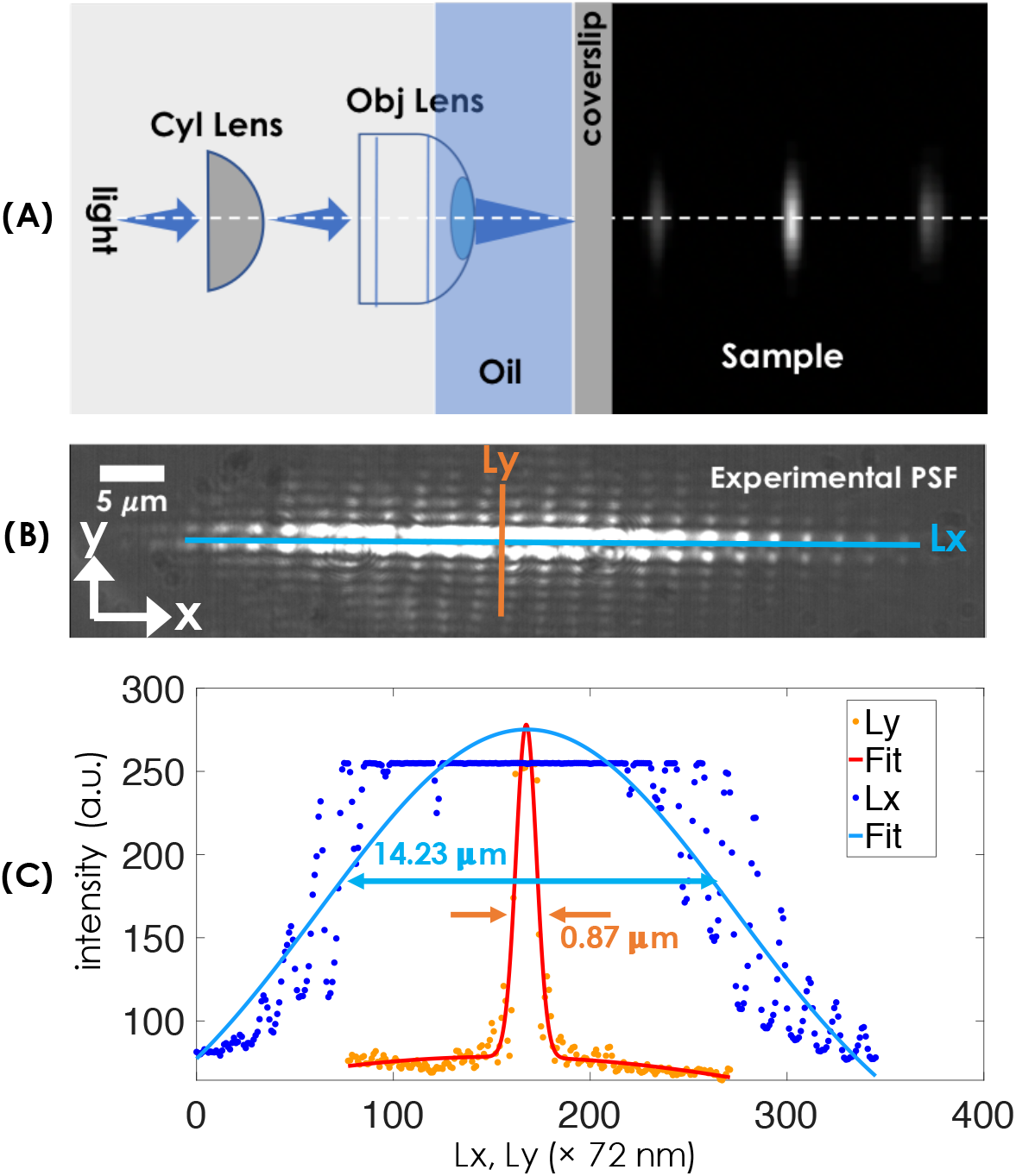
(A) Schematic diagram showing the formation of lightsheet using a combination of cylindrical lens and oil-immersion objective lens in the specimen. (B) The reflected image of lightsheet formed in the cell medium (DMEM) as observed by the camera. (C) Intensity plots across the light sheet (blue and orange line), and the Gaussian fit function show the actual size of lightsheet to be, 0.87 *μm* × 14.24*μm* (*y* × *x*).

Experimentally, the travel time (*t*) can be calculated from the number of frames (of the recorded video) between the initial position (particle undergoing Brownian motion) to the final position (trap-center) using a high-speed camera (CMOS camera). From the video, several beads are tracked on their way to the trap-center, and the time is computed (from top to bottom dotted red line in Fig. 3). Beads take large time or equivalently more number of frames (represented by blue dots) to reach trap-center at low light compared to intense light (see inset in Fig. 3). This is understandable since low power produces a weak optical trap. Other important parameters include mass (*m* = *ρV*) of the bead can be calculated from the density of bead ~ 2000 *Kg/m*^3^, and its volume (assuming spherical shape, *V* = (4/3)*π*(*d*/2)^3^, where *d* is the diameter). The average time between two consecutive positions of the bead or equivalently the *# frames* (represented by blue dots in the track-plot) is chosen as 9.4 *msec* (camera frame-rate). Knowing that, the viscosity of water at 25°*C* is, *η* = 0.8925 ×10^−3^ *Pa · s*, the average trap stiffness of LOT can be calculated using, *k* = 6*πηr_b_v/t* = 16.82 ×10^−9^/*t pN*/*nm*. A better estimate can be arrived at by taking into account other forces. Fig. 3 shows the trap stiffness (*k*) at varying light intensity, with a maximum of 0.00118 *pN/nm* at 52.5 *mW*. It is immediately evident that lightsheet traps are an order weaker than typical point traps [41] [42]. This is predominantly because light sheet PSF is spread over a larger space (here along a line) than a point, so the intensity is much weaker than that of a typical point-trap.

**FIG. 3:**
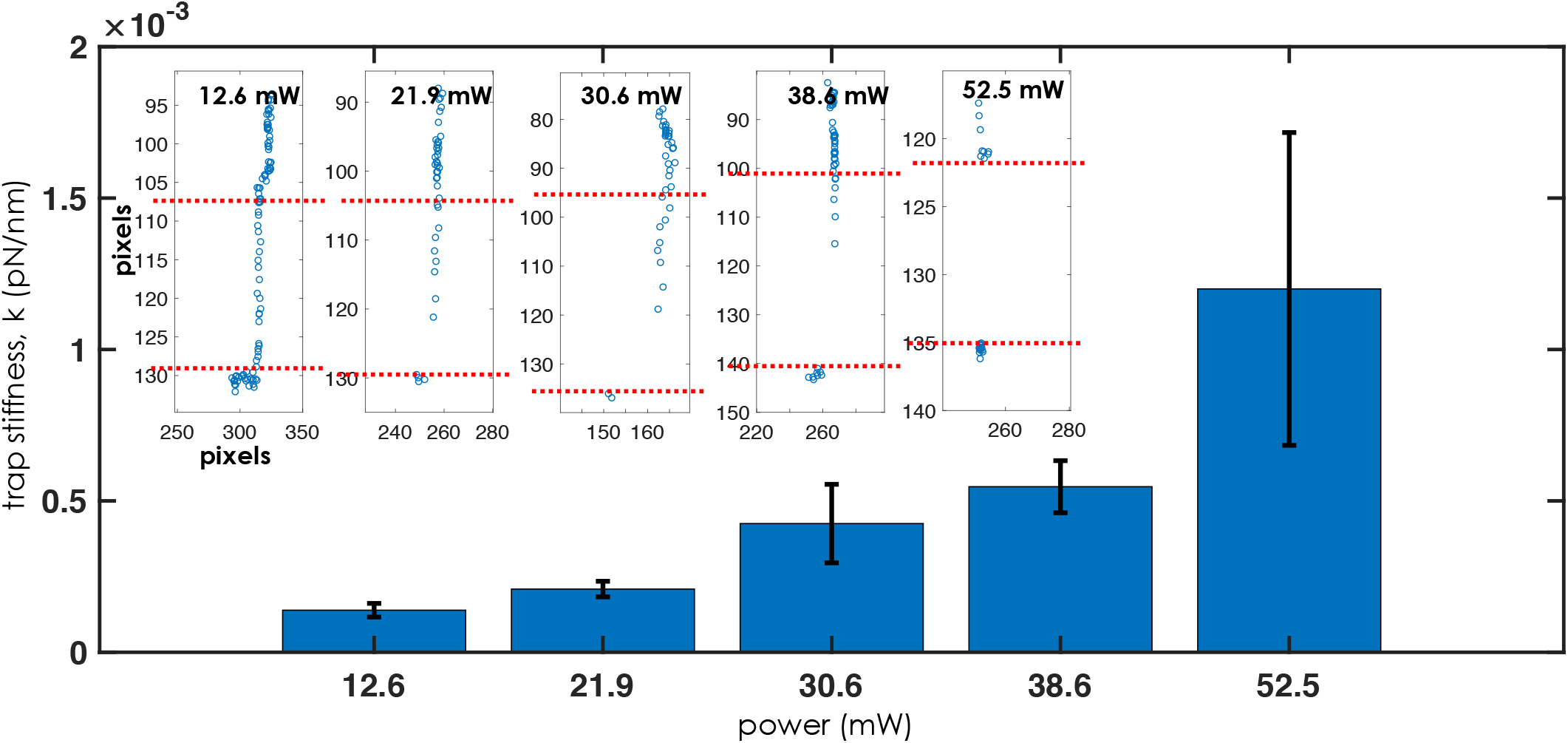
Trap stiffness *k* (*pN*/*nm*) of LOT at varying light intensity, 12.6 – 52.5 *mW*. The insets show the time taken by a single bead to reach the trap-center with increasing intensity, where the number of blue dots indicates the number of frames taken during the trapping process.

To visualize the functioning of LOT system, we used dielectric beads suspended in distilled water as a sample. A drop of the bead solution is dropped on the glass-coverslip, and then it is placed on the oil-dipped 100X objective lens. The light sheet is generated in the solution as shown in Fig. 4. The beads can be seen randomly distributed with two beads lying on the light sheet. At the time *t* = 13.18 *secs*, two beads are trapped, and one bead (marked by violet arrow and circle) is near the light sheet. In the frame (recorded at 13.46 *secs*), the bead is seen trapped by the gradient potential. The next frame (taken at *t* = 15.62 *secs*) shows an approaching random bead (marked by the blue arrow) which is eventually attracted by the potential in the subsequent two frames. The same can be observed for the next bead (marked by a red arrow). Over time a number of beads are arranged on a line (light sheet), as seen from the image taken at 37.60 *sec*. The entire trapping process can be visualized in the **Supplementary Video 1**.

**FIG. 4:**
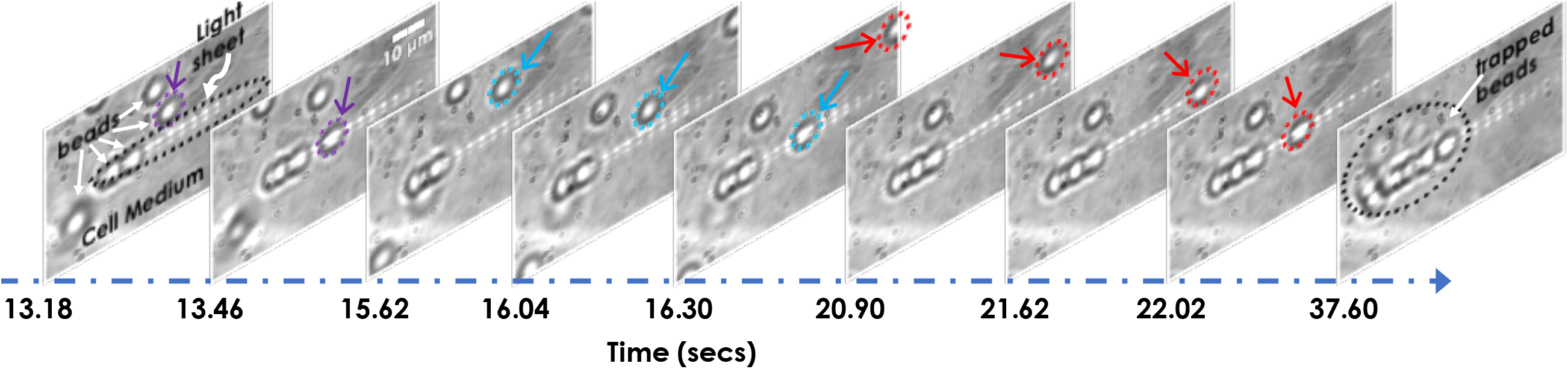
Images taken during sequential trapping of several beads to pattern beads in the form of a chain. All the beads displayed undergoing Brownian motion. Few beads in close proximity to the light sheet are indicated by violet, blue and red circle/arrow that are eventually trapped in the process. The respective timeline is shown below. The entire process is encapsulated in the **Supplementary Video 1**.

Cell-cell interaction (interaction between cell surfaces) plays a critical role during the early development of multicellular organisms. These interactions allow cells to communicate with each other in the microenvironment (culture medium). It is the communication between the cells that control its growth at a healthy rate. Uncontrolled growth is known to occur in cancer. We have considered HeLa cancer cells for the present study. The cells were thawn and grown in a 35 mm disc supplemented with cell medium (DMEM+FBS). To detach them from the surface, the cells were tripsinated followed by centrifugation and resuspension according to sample preparation protocols detailed in **Supplementary Material**. Subsequently, the floating cells (spherical shape) were subjected to lightsheet trap. One-by-one the cells were trapped by the light sheet field and aligned in a line as displayed in Fig. 5 (see, blue, red, and yellow arrow). The corresponding timeline is also indicated. The entire process can be visualized in **Supplementary Video 2**. The results show that the technique can pattern cell in a preferential direction (along a line) that can help understand cell-cell interaction both at a single cell level and in an ensemble. The next logical step is to understand patterned cell growth.

**FIG. 5:**
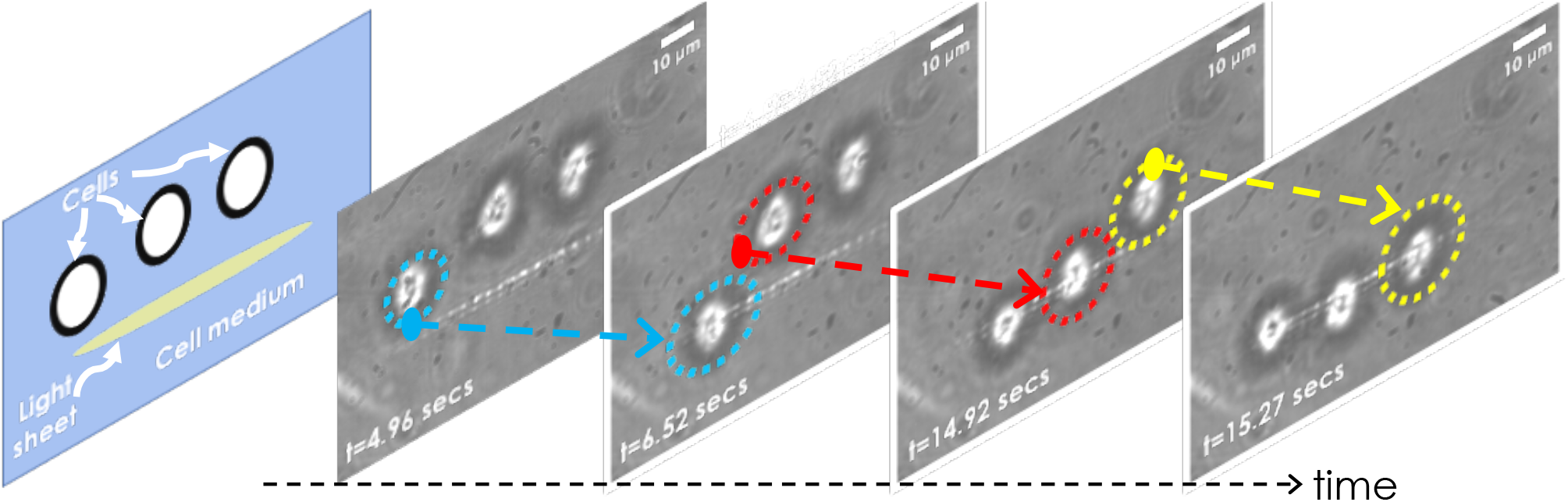
LOT technique is used to trap HeLa cells suspended in cell medium (DMEM). The cells were trapped and manipulation one-by-one, and aligned in a line. See **Supplementary Video 2**.

We propose and demonstrate the first lightsheet tweezer for trapping dielectric particles and live HeLa cells. Unlike traditional point-trap tweezers, LOT traps are relatively weak but strong enough to align particles in a line that exhibit one-degree of freedom along the light sheet axis. LOT is used to demonstrate the trapping of dielectric beads suspended in distilled water and HeLa cells in DMEM. From the time-travel of particle (bead) to the epicenter of trap, the stiffness of 1D potential is determined by calculating time using a high-speed CMOS camera. We showed maximum trapping of 8 beads and 3 HeLa cells in a light sheet. LOT is highly scalable and can be used to trap a few to several particles. Further progress of LOT may lead to 2D planar traps using a relatively large uniform light sheet that can be obtained by using a low NA objective and a high laser power. LOT may find immediate application in colloidal physics, particle trapping, atomic physics, and patterned cell growth in cell biophysics.

## Acknowledgements

The authors acknowledge financial support from parent institute (Indian Institute of Science, Bangalore, India). PPM conceived the idea. AS, JB, PJ, NB, PPM carried out the experiment. AS, PJ, NB prepared the sample. PPM wrote the paper by taking inputs from all the authors.

## Data Availability

The data that support the findings of this study (including the 3D volume of the specimens) are available from the corresponding author upon request.

