## Supplementary Material for "Lightsheet Optical Tweezer (LOT) for Optical Manipulation of Microscopic Particles and Live Cells"

Actual LOT System:

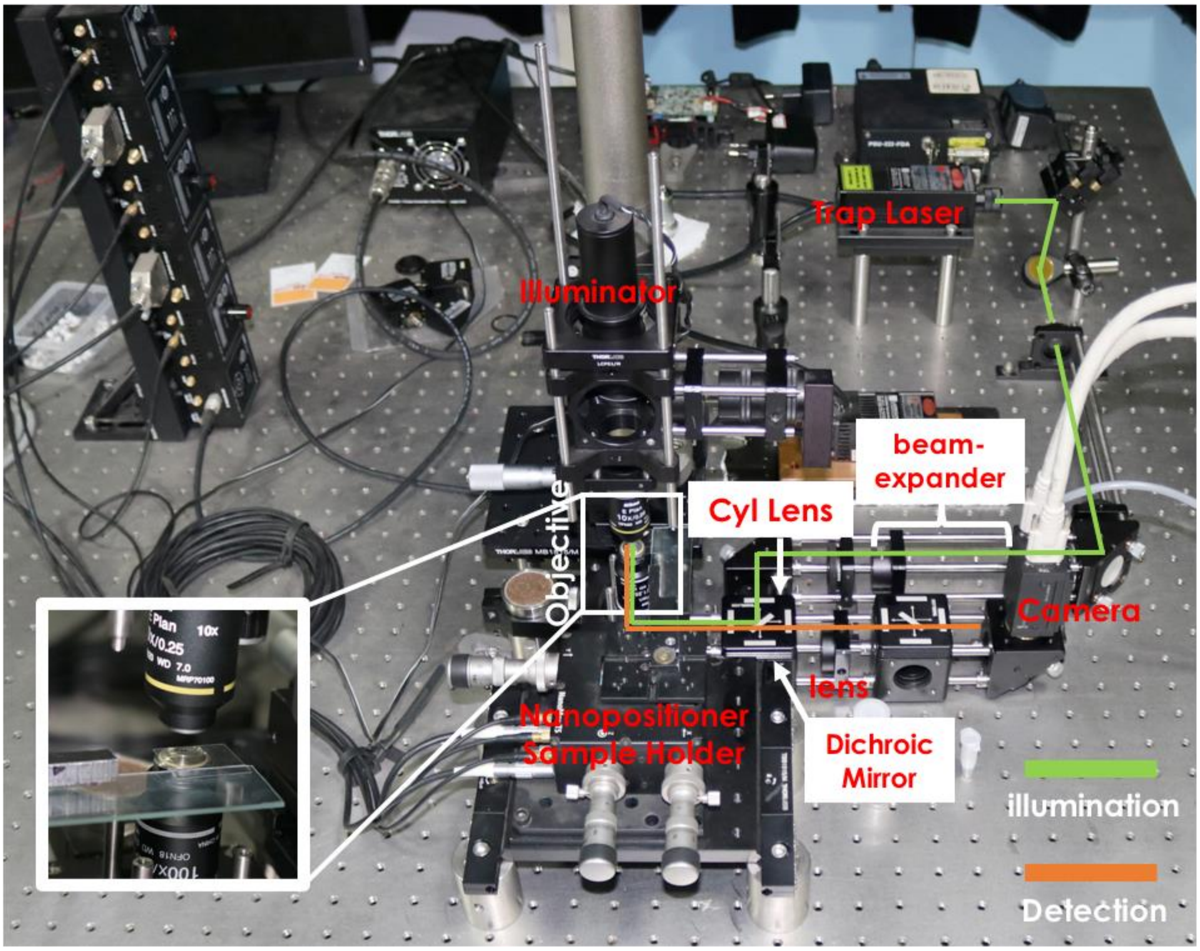


**Fig. S1:** The actual experimental setup of LOT system. The illumination and the detection arms are indicated by green and orange lines, respectively. Most of the key components are also indicated.

The optical setup for lightsheet optical tweezer (LOT) is shown in Fig. S1. A high power laser (of maximum power 500 mW) with a trap wavelength of 1064 nm is used to trap particles and live cells. The light is directed to the cylindrical lens through a series of IR mirrors (using both adjustable and fixed mirror holders). The beam is aligned to the center of 25 mm optics using components purchased from Thorlabs. The cylindrical lens (of focal length = 150 mm) is positioned such that it focus the beam at the back aperture of objective lens. This results in the formation of diffraction limited tightly-focused light-sheet at the working distance of the objective lens. We used a high NA objective lens for tight focusing of the beam. This results in a line focus rather than a sheet focus. The samples (beads and live Hela Cells) are dropped on a coverslip attached to a glass slide which is placed on a nano-positioner for fine movement of the sample stage. Trapping of both beads and cells are successfully achieved as shown in **Supplementary Video 1 and 2**. A top white light illuminator is used to visualize the beads in the camera. On its way to the detection, the trap laser light is filtered out by the dichroic mirror. Due to about 80% transmission efficiency of the filter, we are able to see part of light sheet as well. The dichroic mirror in combination with a shortpass filter prevents backscattered light from the 1064 nm laser from saturating the CMOS sensor.

**Sample Preparation:**

**Beads Sample:** We have purchased non-Functionalized Fused Silica Beads in Deionized Water from Thorlabs. Subsequently, the beads (of size ~ 2.06 μm) are diluted to one-fourth of the original concentration of 2 gm/ml.

**HeLa Cells and maintenance:** The HeLa cells were cultured in complete Dulbecco's modified minimal Eagle's medium (DMEM) (Gibco, Thermo Fisher Scientific) supplemented with 10% FBS( Gibco, Thermo Fisher Scientific) and 1% penicillin –streptomycin solution (Gibco, Thermo Fisher Scientific) at 37°C and 5% $CO_2$ ($CO_2$-incubator, Thermo Scientific). Hemocytometer is used to count cells after every passage and approximately 100,000 cell count was maintained. The cells were passaged in every 2 -3 days to maintain healthy cell lines. After two passages, the cells were tripsinated using 3.7% Tripsine which is followed by 4 min incubation. The cells were then pipetted out to break-down cell-clusters and centrifuged at 3000 rpm. The supernatant is then pipetted out, and the cell-pallet were resuspended in DMEM. These cells were kept for 5 minutes in the incubator before carrying out trapping experiments.
